# Structure-guided engineering of a fast genetically encoded sensor for real-time H_2_O_2_ monitoring

**DOI:** 10.1101/2024.01.31.578117

**Authors:** Justin Daho Lee, Woojin Won, Kandace Kimball, Yihan Wang, Fred Yeboah, Kira M. Evitts, Carlie Neiswanger, Selena Schattauer, Michael Rappleye, Samantha B Bremner, Changho Chun, Netta Smith, David L. Mack, Jessica E. Young, C. Justin Lee, Charles Chavkin, Andre Berndt

**Affiliations:** Molecular Engineering and Sciences Institute, University of Washington, Seattle, WA, USA; Department of Bioengineering, University of Washington, Seattle, WA, USA; Institute for Stem Cell and Regenerative Medicine, University of Washington, Seattle, WA, USA; Center for Neuroscience of Addiction, Pain and Emotion, University of Washington, Seattle, WA, USA; Center for Cognition and Sociality, Institute for Basic Science, Daejeon, Republic of Korea; Department of Laboratory Medicine and Pathology, University of Washington, Seattle, WA, USA; Department of Rehabilitation Medicine, University of Washington, Seattle, WA, USA

**Author notes:** now at: Institute of Pharmacology and Toxicology, University of Zurich, Zurich, Switzerland (Michael Rappleye). Co-senior authors.

## Abstract

Hydrogen Peroxide (H_2_O_2_) is a central oxidant in redox biology due to its pleiotropic role in physiology and pathology. However, real-time monitoring of H_2_O_2_ in living cells and tissues remains a challenge. We address this gap with the development of an optogenetic hydRogen perOxide Sensor (oROS), leveraging the bacterial peroxide binding domain OxyR. Previously engineered OxyR-based fluorescent peroxide sensors lack the necessary sensitivity or response speed for effective real-time monitoring. By structurally redesigning the fusion of Escherichia coli (E. coli) ecOxyR with a circularly permutated green fluorescent protein (cpGFP), we created a novel, green-fluorescent peroxide sensor oROS-G. oROS-G exhibits high sensitivity and fast on-and-off kinetics, ideal for monitoring intracellular H_2_O_2_ dynamics. We successfully tracked real-time transient and steady-state H_2_O_2_ levels in diverse biological systems, including human stem cell-derived neurons and cardiomyocytes, primary neurons and astrocytes, and mouse neurons and astrocytes in ex vivo brain slices. These applications demonstrate oROS’s capabilities to monitor H_2_O_2_ as a secondary response to pharmacologically induced oxidative stress, G-protein coupled receptor (GPCR)-induced cell signaling, and when adapting to varying metabolic stress. We showcased the increased oxidative stress in astrocytes via Aβ-putriscine-MAOB axis, highlighting the sensor’s relevance in validating neurodegenerative disease models. oROS is a versatile tool, offering a window into the dynamic landscape of H_2_O_2_ signaling. This advancement paves the way for a deeper understanding of redox physiology, with significant implications for diseases associated with oxidative stress, such as cancer, neurodegenerative disorders, and cardiovascular diseases.

## Introduction

Endogenous Reactive Oxygen Species (ROS) are indispensable components of aerobic metabolism, which hallmarks the rise of complex life.^1,2^ Due to their damaging impact on biological macromolecules, redox homeostasis is tightly regulated in most aerobic systems, and high-level accumulation of ROS is often viewed as a pathogenic marker in degenerative diseases (e.g. Alzheimer’s disease, Duchenne Muscular Dystrophy), tumorigenesis, and inflammation.^3–6^ Furthermore, an increasing number of studies report the role of low-level ROS as physiologic mediator in normal cellular signaling processes^7–9^. Specifically, H_2_O_2_ is a key redox signaling molecule, owing to its relative stability and ability to modify cysteine residues in proteins, enabling selective downstream signaling.^10^ On the other hand, excessive H_2_O_2_ is a common pathological marker affecting phenotypic and disease progression in various cell types.^11–13^ Nevertheless, limited analytic tools to spatiotemporally monitor specific oxidants *in situ* with precision have been a bottleneck to deciphering their specific role in physiology and the cause and effect of their imbalance.^14,15^ Thus, methods to interrogate the role of H_2_O_2_ would be broadly applicable to the study of redox biology and medicine.^15^

Most synthetic ROS-sensitive dyes are unsuited for these considerations because of their short working time window, low sensitivity, and low specificity.^16^ Protein-based peroxide sensors have been engineered to overcome these shortcomings. For example, the roGFP sensor family fuses roGFP, a redox-sensitive green fluorescent protein variant, to H_2_O_2_-specific enzymes like Orp1 (thiol peroxidase), or Tsa2 (typical 2-Cys peroxiredoxin) from yeast to achieve peroxide-specific roGFP fluorescence changes via redox relay.^17,18^ The HyPer sensor family is based on the direct fusion of circularly permuted fluorescent protein (cpFP) to the regulatory domain of bacterial peroxide sensor protein OxyR for conformational coupling that leads to H_2_O_2_-specific fluorescence change.^19–24^ Most HyPer sensors use ecOxyR (*Escherichia coli* OxyR), the most extensively studied OxyR variant, as their sensing domain. However, existing ecOxyR-based peroxide sensors exhibit low sensitivity and slow oxidation kinetics (seconds under saturation conditions)^21,22,25^, while studies reported peroxide-dependent oxidation of ecOxyR at a sub-second scale^26^. We hypothesized that the discrepancy stems from the disruption of structural flexibility in the sensors. Through a series of structure-guided engineering steps, we developed oROS-G (optogenetic hydRogen perOxide Sensor, Green), a green fluorescent protein (GFP, excitation: 488 nm, emission: 515 nm) and an ecOxyR-based peroxide sensor that exhibits exceptional sensitivity and kinetics enabling the visualization of peroxide diffusion. We also engineered oROS-Gr, a ratiometric variant of oROS-G by fusing it with mCherry, which allows measurement of the precise sensor oxidation state by normalizing sensor fluorescence intensity for the expression level. Here, we present diverse use cases of oROS sensors to monitor both steady-state and transient H_2_O_2_ levels in various model systems. Additionally, we showed how oROS can detect varying H_2_O_2_ levels in astrocytes in the context of Alzheimer’s disease models and assessed the efficacy of a drug in reducing aberrant peroxide levels. Also, we investigated how different glucose levels can result in different intracellular oxidative environments in conjunction with mitochondrial respiratory depression. Lastly, we showed morphine-dependent peroxide generation in µ-opioid receptor-expressing neurons in the brain of mice *ex vivo*.

## Result

### Structure-guided engineering strategies for ecOxyR-based H_2_O_2_ sensor with improved sensitivity and kinetics

OxyR is a bacterial H_2_O_2_ sensor protein that regulates the transcription of antioxidative genes in response to low-level cellular H_2_O_2_. The specificity of OxyR for H_2_O_2_ stems from its unique H_2_O_2_ binding pocket^27^. Previous studies have shown that binding H_2_O_2_ leads to an intermediate state that facilitates the disulfide bridging of two conserved cysteine residues (C199-C208), which triggers the transition into the oxidized conformational state of OxyR. Due to its unique characteristic as an H_2_O_2_ sensor with low scavenging capacity^27^, OxyR is attractive for building a protein-based H_2_O_2_ reporter. Nevertheless, the slow kinetics and low sensitivity of existing ecOxyR sensors^19,21–23,25^ deviate from the reported ecOxyR kinetics, prompting us to revisit the sensor design. **[Fig. 1A, Supp. Fig. 1A]** OxyR-based peroxide sensors have circular permuted fluorescent proteins (cpFP) within the loop between residues C199 and C208. However, the crystal structure of oxidized ecOxyR [PDB:1I6A] predicted an evident peak of B-factor **[Fig. 1B]** indicating this loop region is more flexible than its surroundings. We hypothesized that inserting the bulky cpFP there (e.g. in HyPer sensors) could diminish sensing performance by possibly increasing the conformational entropy of the intermediate state that brings C199 and C208 into proximity^27^. We performed pairwise residue distance analysis between oxidized and reduced ecOxyR structures and found that the region between residues 209-220 goes through noticeable peroxide-dependent conformational change. **[Supp. Fig. 1B]** Therefore, we tested alternative cpGFP insertion within this region. The functional screening for oROS sensors was performed in Human Embryonic Kidney (HEK293) cells to ensure compatibility with other mammalian host systems. cpGFP insertion between residue 211 and 212 elicited a robust response (97.55% in ΔF/Fo; confidence interval 95% (ci) = [96.6, 98.52]) to 300 µM extracellular peroxide, which has been reported to induce full oxidation of OxyR-based sensors^22^ **[Fig. 1B, C].** The 211-212 variant responded immediately (in 25-75% sensor saturation response kinetics, 1.06s; ci = [1.05, 1.07]) which was not observed in other ecOxyR-based sensors^19,21,22,25^. Moreover, the variant showed improved response amplitudes (20.41% in ΔF/Fo; ci = [19.62, 21.17]) to low peroxide levels (10µM) compared to HyperRed (2.8% in ΔF/Fo; ci = [2.61, 3.0]), which inserted the red fluorescent protein cpmApple between OxyR positions 205 and 206 **[Fig. 1D].** Next, based on the guiding principles learned from engineering of the calcium indicator GCaMP5^28^, we introduced large and apolar amino acid tyrosine at the residue sites putatively proximal to the cpGFP predicted opening to reduce solvent access. We found the E215Y mutation increased response amplitude (ΔF/Fo) by 2.1-fold at full oxidation (ci = [1.99, 2.26]) and we named this variant oROS-G **[Fig. 1E, Supp. Fig. 1C]**.

**Figure 1.**
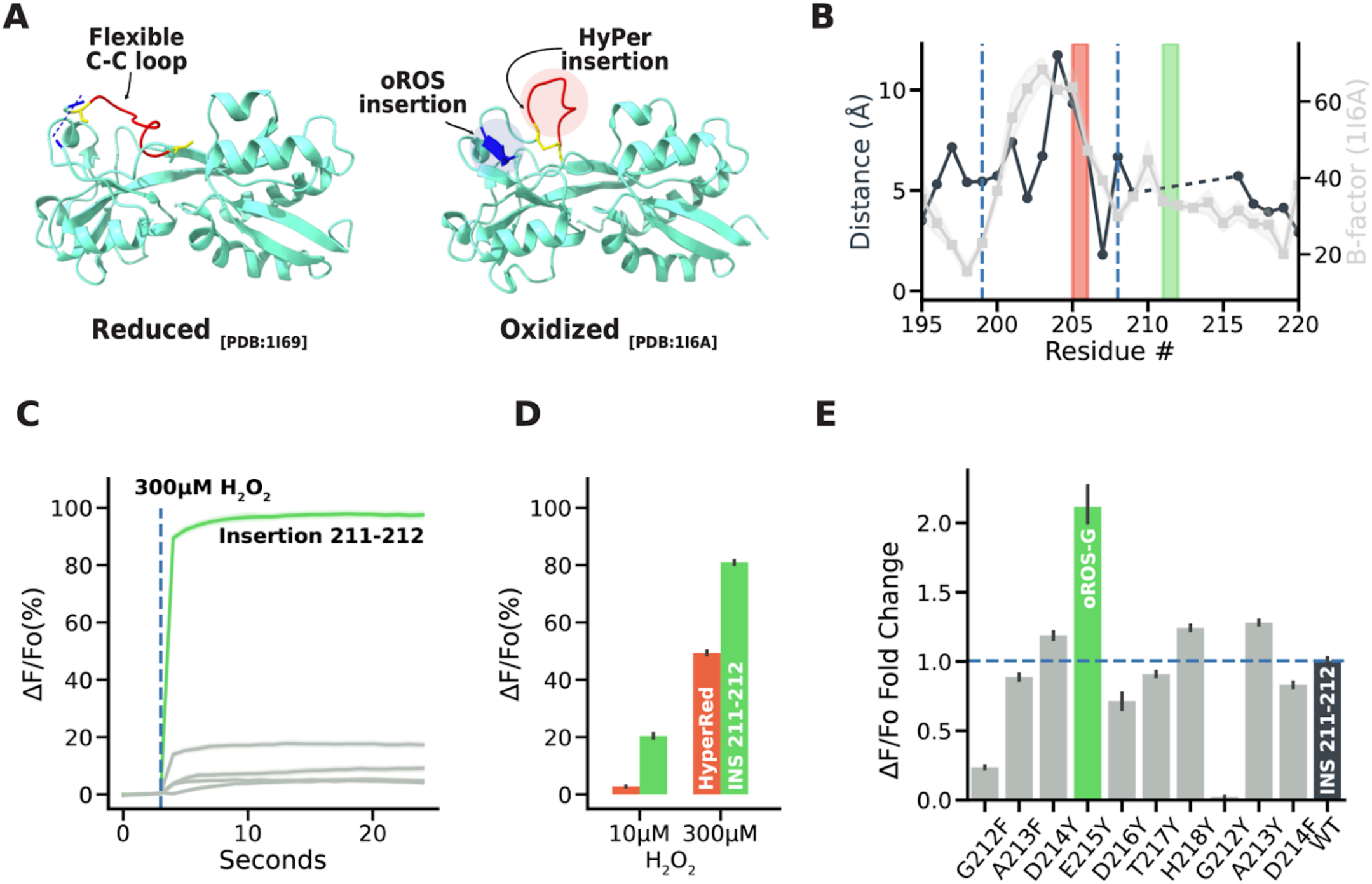
Structure-guided engineering strategies for ecOxyR-based H_2_O_2_ sensor **A-B** Structure-guided hypothesis of oROS sensor design. **A** Crystal structure of reduced and oxidized forms of Regulatory Domain (RD) of Escherichia coli OxyR. Cystine-cystine pair labeled in yellow. Red indicates the fluorescent protein insertion loop for HyPer sensors, and Blue indicates the newly identified fluorescent protein insertion site for oROS sensors. **B** Overlay of B-factor of oxidized OxyR structure and residue-to-residue distance plots for a zoomed-in view of the putative region with high conformational change between oxidized and reduced form of ecOxyR. The red and green boxes indicate the insertion sites of HyPerRed and oROS-G, respectively. The insertion site proposed for oROS-G is outside of the loop between C199 and C208 (gray line). To maximize the flexibility of the loop. **C-E** Screening of oROG-G sensor variants. All the sensor variants were expressed and screened on HEK293 cells (n>100 cells per condition/variant). **C** fluorescence change (ΔF/Fo) in response to extracellular H_2_O_2_ (300µM) stimulation on variants with cpGFP insertion to newly identified oROS insertion region. Insertion 211-212, a variant with exceptional response kinetic and dynamic range was identified. **D** Maximum fluorescence change (ΔF/Fo) of Insertion 211-212 and HyPerRed in response to high (300µM) and low(10µM) extracellular H_2_O_2._ **E** Maximum fluorescence change (ΔF/Fo) of site-directed mutagenesis variants predicted to reduce water access for cpGFP. Mutation E215Y on Insertion 211-212 variant led to engineering oROS-G. **Descriptive statistics:** Error bars and bands represent the bootstrap confidence interval (95%) of the central tendency of values using the Seaborn (0.11.2) statistical plotting package. Cell-of-interests were collected from 3 biological replicates unless noted otherwise.

### Characterization of ultrasensitive and fast peroxide sensor, oROS-G

We first characterized the fluorescence response of the oROS-G sensor in HEK293 cells in response to exogenously or endogenously sourced H_2_O_2_. Direct application of exogenous H_2_O_2_ increases intracellular H_2_O_2_ by diffusion across the plasma membrane through specific aquaporins, which creates an extracellular-to-intracellular gradient of H_2_O_2_ ^29–31^. Under these conditions, the intracellular concentration of H O is reported to be about 2 or 3 magnitudes lower than that of extracellular H_2_O_2_ ^22,32^. On the other hand, the pharmacological agent menadione produces H_2_O_2_ intracellularly through various redox cycling mechanisms^33^. The signal amplitude of oROS-G (192.34% in ΔF/Fo; ci = [190.45, 194.23]) at saturation (300 uM H_2_O_2_) was ≈2-fold that of HyPerRed (97.74% in ΔF/Fo; ci = [96.52, 99.06]), Improved sensitivity of oROS-G yielded a ≈7.08 times larger response at low-level peroxide stimulation. (oROS-G: 116.22% in ΔF/Fo; ci = [110.85, 121.73] vs HyPerRed: 16.45% in ΔF/Fo; ci = [15.98, 16.95]) [**Fig. 2A**] oROS-G also exhibited significant improvement in on-kinetics compared to HyPerRed with ≈38 times faster 25-75% ΔF/Fo kinetics **[Fig. 2B]** Intriguingly, the fast oxidation kinetics of the oROS-G sensor captured the H_2_O_2_ diffusion across the imaging field of view from the media mixing, in contrast to HyPerRed which exhibited uniform population response. [**Supp. Fig. 2A, B]** Further analysis revealed the speed of peroxide diffusion during media mixing to be ≈824µm/s. **[Supp. Fig. 2C, D]** The speed of peroxide travel slows down to ≈100µm/s after passing the cell plasma membrane during intracellular diffusion. This potentially represents peroxide travel becoming rate limited by aquaporin-driven passive transmembrane diffusion^30,34^ **[Supp. Fig. 2E, F].** Taken together, visualization of bolus H_2_O_2_ introduction using oROS-G was only rate-limited by H_2_O_2_ travel speed and transmembrane transport rate, allowing real-time observation of intracellular peroxide diffusion in mammalian cells. Thus, oROS-G can be a vital tool for expanding our understanding of the dynamic topological and temporal landscape of peroxide in biological systems.

**Figure 2.**
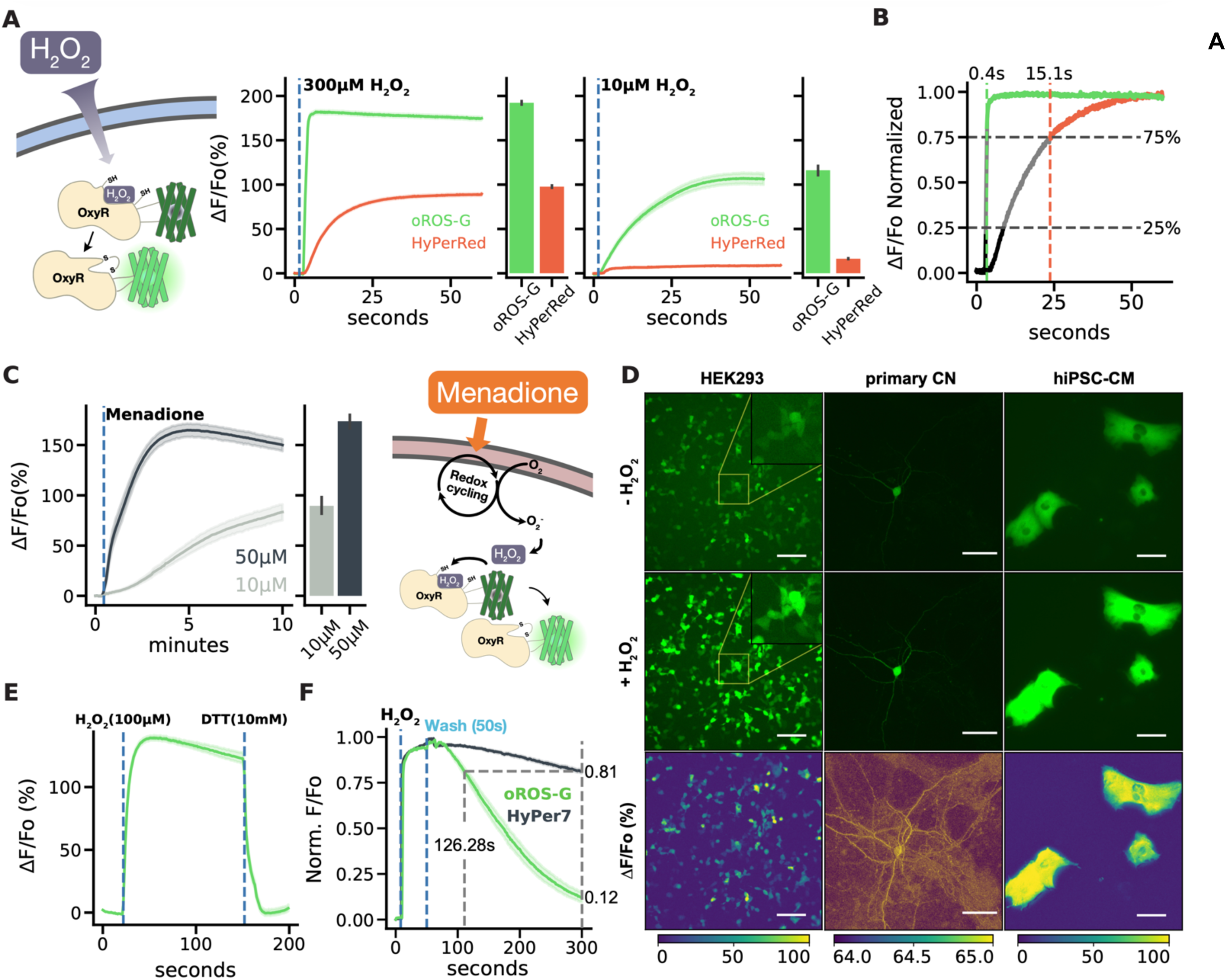
Characterization of ultrasensitive and fast peroxide sensor, oROS-G. **A *Left*** Fluorescence responses of oROS-G and HyPerRed to 300µM exogenous H_2_O_2_. The barplot represents the mean of the maximum fluorescent response of cells. ***Right*** Fluorescence responses of oROS-G and HyPerRed to 10µM exogenous H_2_O_2_. The barplot represents the mean of the maximum fluorescent response of cells. **B** On-kinetic analysis of oROS-G and HyPerRed. Representative trace of 300µM peroxide-induced saturation of both sensors with normalized ΔF/Fo. Vertical dotted lines represent 75% sensor activation, labeled with 25-75% activation completion time. 100% indicates full sensor saturation. **C** Fluorescence response of oROS-G to high (50µM) or low (10µM) menadione (n>100 cells per condition). The sensors were expressed in HEK293. The barplot represents the mean of the maximum fluorescent response of cells. **D** Expression of oROS-G in HEK293 (Scale bar: 100µM), primary rat cortical neuron (Scale bar: 50µM), and hiPSC-derived cardiomyocyte (Scale bar: 100µM), and their responses to 300µM H_2_O_2_ stimulation. **E** Induced reduction of oROS-G. HEK293s expressing oROS-G were first stimulated with 100µM H_2_O_2_ and then 10 mM Dithiothreitol (DTT), a reducing agent, shortly after (n>100 cells). **F** Representative reduction kinetics of oROS-G and HyPer7 after 100µM H_2_O_2_ stimulation followed by media wash (n>100 cells per sensor). **Descriptive statistics:** Error bars and bands represent the bootstrap confidence interval (95%) of the central tendency of values using the Seaborn (0.11.2) statistical plotting package. Cell-of-interests were collected from 3 biological replicates unless noted otherwise.

In HEK293 cells, oROS-G also acutely responded to 10µM and 50µM menadione in a dose-dependent manner. (ΔF/Fo, 10µM: 89.56 %; ci = [81.79, 97.57], 50µM: 173.68%; ci = [166.81, 180.35]) **[Fig. 2C]** The result was consistent in human primary astrocytes, **[Supp. Fig 3A]** highlighting the potential robustness of oROS-G expression and functionality in broader biological host systems. In addition, we confirmed the robust expression and function of oROS-G in rat cortical neurons and human-induced pluripotent stem cell-derived cardiomyocytes (hiPSC-CMs) **[Fig. 2D].**

Next, we confirmed that oROS-G is a fully reversible sensor by directly reducing it using 10 mM Dithiothreitol (DTT) **[Fig. 2E]** or media washout of H_2_O_2_ **[Supp. Fig. 2B]**. Here, we noticed that the endogenous reduction kinetics of the sensor in mammalian cells was faster than other OxyR-based sensors^22,24^. For example, both HyPerRed and HyPer7 took 20∼30 minutes for them to return to baseline after the sensor saturation. HyPer7 is the newest green iteration of the HyPer sensor family that was engineered by swapping the sensing domain with a different OxyR domain from *Neisseria meningitidis* (nmOxyR)^24^ with fluorescent reporter insertion contained to the C-C loop region. oROS-G reached ≈90% reduction from its maximum saturation in 4.17 minutes, whereas HyPer7 only achieved about ≈20% reduction from its full saturation in the same duration, consistent with the previous report. (HyPer7: 0.81; ci = [0.8, 0.82], oROSG: 0.12; ci = [0.1, 0.15]) oROS-G showed 2.63 times faster decay kinetics than HyPer7 based on approximation with reduction time to 85% of saturation, making oROS-G a more compelling candidate for measurement of peroxide transient rise and decay of intracellular peroxide species **[Fig. 2F, Supp. Fig. 3C].** Lastly, we created a C199S mutant of oROS-G to show that the fluorescence response was specific to peroxide-induced disulfide bridging of C199-C208, which is consistent with other OxyR-based peroxide sensors. **[Supp. Fig. 3D]**

### Monitoring the effect of antioxidants on intracellular peroxide level in Alzheimer’s model

Next, we explored using oROS-G in the context of antioxidants that target intracellular peroxides. N-acetyl-cystine (NAC) is a cysteine prodrug widely used as a classical “antioxidant”. Although its detailed mechanism of action as not been established, recent studies highlight its antioxidative role via the production of low-level sulfane sulfur species^35,36^. Using oROS-G, we measured the effect of NAC-dependent catabolism on cellular H_2_O_2_ levels in real-time. With a 1-hour preincubation of 1 mM NAC, oROS-G expressed in HEK293 showed a 73% diminished response to exogenous 10 µM peroxide exposure **[Supp. Fig. 4A]**. Similarly, we incubated oROS-G expressing HEK293 cells to either NAC (10 mM) or Vehicle (DMSO) for 20 minutes before 10 µM menadione exposure. NAC significantly attenuated the response by 72 percent **[Supp. Fig. 4B]**. We next examined the ability of the oROS-G sensor in primary cultured astrocytes to quantitatively detect endogenous H_2_O_2_ levels to assess the H_2_O_2_-scavenging effects of molecules. Initially, we expressed oROS-G in the astrocytes and tested H_2_O_2_ concentration-dependent functionality **[Supp. Fig. 4C]**. We observed an increase of 21.27 ± 5.3% and 57.74 ± 9.4% in fluorescence with 10 and 100 μM H_2_O_2_, respectively **[Supp. Fig. 4D-F]**, indicating a functional response of the oROS-G sensor in primary cultured astrocytes. Previously, we have shown that aberrant H_2_O_2_ production by reactive astrocytes, a pathological form of astrocytes, is a contributing factor to Alzheimer’s disease (AD) pathology^11,37^. In detail, upregulated monoamine oxidase B (MAOB) in reactive astrocytes produces H_2_O_2_ by breaking down oligomerized amyloid beta (oAβ)-metabolites, such as putrescine, leading to pathological H_2_O_2_ generation^38,39^ **[Fig. 3A].** Therefore, we tested whether we could monitor aberrant endogenous H_2_O_2_ production in oROS-G transfected astrocytes treated with oAβ (5 μM) or putrescine (180 μM). Over a 40-hour continuous recording **[Fig. 3B]**, we observed a significant increase in oAβ-induced oROS-G fluorescence, indicating a notable rise in H_2_O_2._ **[Fig. 3C, D].** Conversely, the application of KDS2010 (1 μM), a selective MAOB inhibitor, and sodium pyruvate (1 mM), a potential H_2_O_2_ scavenger, showed a smaller increase in H_2_O_2_ levels. Additionally, incubation with putrescine, a pre-substrate of MAOB, also significantly increased oROS-G sensor fluorescence **[Fig. 3E, F]**. However, this H_2_O_2_ elevation was significantly reduced by KDS2010 and partially reduced by sodium pyruvate. Taken together, these results suggest that the oROS-G sensor in primary cultured astrocytes is a reliable tool for monitoring endogenous H_2_O_2_ production under AD-like conditions and evaluating the efficacy of potential H_2_O_2_-scavenging compounds.

**Figure 3.**
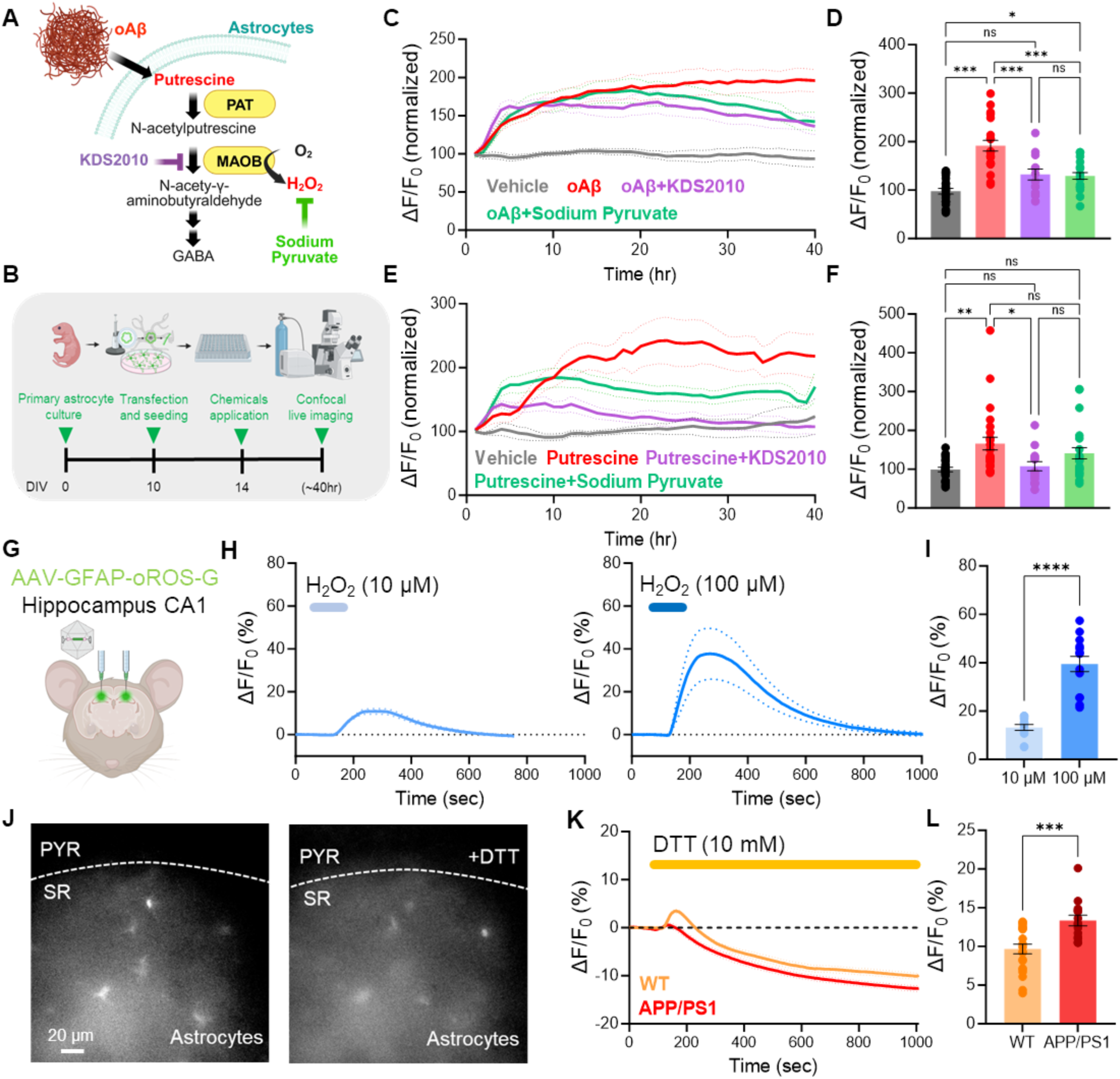
Monitoring the effect of antioxidants on intracellular peroxide level in an Alzheimer’s model. **A** Schematic illustration of the experimental timeline showing oROS-G transfected primary astrocytes seeded in a 96-well plate, administered with 5 µM oligomerized Amyloid beta (oAβ), 180 µM putrescine, 1 µM KDS2010 (selective and reversible MAOB inhibitor), and 1mM sodium pyruvate (H_2_O_2_ scavenger) followed by 40-hour of confocal live imaging to monitor intracellular H_2_O_2_ and antioxidant effects. **B** Schematic diagram of the oAβ and putrescine induced H_2_O_2_ production. **C** 40-hour monitoring of oAβ-induced H_2_O_2_ production and antioxidant effects with KDS2010 and sodium pyruvate (Vehicle: n=22, oAβ: n=22, oAβ+Sodium Pyruvate: n=23, oAβ+KDS2010: n=13). **D** Summary bar graph representing fluorescence values normalized to the baseline, measured during the first hour. **E** Monitoring of putrescine-induced H_2_O_2_ production and antioxidant effects with KDS2010 and sodium pyruvate (Vehicle: n=24, Put.: n=25, Put.+Sodium Pyruvate: n=22, Put.+KDS2010: n=14). **F** Summary bar graph representing fluorescence values normalized to the baseline. **G** Schematic illustration of the bilateral virus injection of experimental design. **H** Fluorescence response of oROS-G to 10 µM (n=13) or 100 µM H_2_O_2_ (n=14) in astrocytes of hippocampal tissue. **I** H_2_O_2_ dose-dependent summary bar graph. **J** Representative images captured with a CMOS camera illustrating the expression of oROS-G in astrocytes located in the striatum radiatum (SR) of the hippocampal CA1 region, in coronal brain slices ex vivo. oROS-G virus, driven by the GFAP promoter, is selectively expressed in astrocytes, with no expression in pyramidal neurons (PYR). **K** Fluorescence response of oROS-G to 10 mM (DTT) in astrocytes of hippocampal tissue (WT: n=22, APP/PS1 n=14). **L** Summary bar graph representing the difference between baseline fluorescence and saturation values after DTT administration. **Descriptive statistics:** Data are presented as mean ± SEM. **Inferential statistics:** D, E - One-way ANOVA with Tukey’s multiple comparison test, I,J - Unpaired t-test, two-tailed *P < 0.05, **P < 0.01, ***P < 0.001.

Then we asked whether we could measure the endogenous H_2_O_2_ levels in mouse brains. To test this idea, we bilaterally injected the AAV5-GFAP104-oROS-G virus into the CA1 hippocampus of APP/PS1 mice^39,40^, a well-known AD model, to overexpress oROS-G sensor specifically in the astrocytes **[Fig. 3G].** Two weeks post-injection, we prepared brain slices and tested the H_2_O_2_ concentration-dependent functionality of the sensor. Again, to test the functionality of the sensor, we applied 10 and 100 μM H_2_O_2_ through bath application. We found an increase of 13.24 ± 1.2% and 39.46 ± 3.12% in fluorescence with 10 and 100 μM H_2_O_2_, respectively **[Fig. 3H, I].** Like *in vitro*, the oROS-G sensor functions effectively in astrocytes *ex vivo*. Next, we examined the capability to measure elevated H_2_O_2_ levels in astrocytes of APP/PS1 mice. We hypothesized that treatment with DTT would unmask the portion activated by astrocytic H_2_O_2_. Following DTT (10 mM) administration, we observed a reduction in fluorescence below the baseline levels. Notably, we demonstrated that APP/PS1 mice exhibited a greater reduction compared to wild-type, suggesting a potential method for measuring endogenous H_2_O_2_ levels **[Fig. 3J-L].** Taken together, these results demonstrate that the oROS-G sensor functions effectively *ex vivo*, presenting a potential method for measuring endogenous H_2_O_2_ levels and investigating the antioxidant capacity of various molecules.

### oROS-Gr for long-term and non-continuous monitoring of intracellular H_2_O_2_

Most ratiometric sensors designed for peroxide response are based on dual-excitation of the green fluorescent sensor proteins at 405 nm and 488 nm^41,42^. Although this sensor type allows flexibility in multi-color optogenetic experiments, illumination at 405 nm could contributes to oxidative stress in mammalian cells.^43^ We created oROS-Gr, by fusing mCherry to oROS-G, creating an equimolar reference point inert to H_2_O_2_. Flow cytometry analysis confirmed a strong linear correlation between green (Em. 510 nm) and red (Em. 605 nm) emission intensity of oROS-Gr expressed in HEK293 cells (n = 16,326) **[Fig. 4A]**. Thus, oROS-Gr can be used for long-term and non-continuous monitoring by calculating the green-to-red light emission ratio independent of sensor expression levels. Upon exogenous H_2_O_2_ stimulation, the oROS-Gr ratio (Em. 510/605) showed a dose-dependent response in HEK293 cells. (1 µM: 0.06 (n = 327); ci = [0.06, 0.06], 10 µM: 0.11 (n = 306); ci = [0.1, 0.11], 25 µM: 0.14 (n = 405); ci = [0.14, 0.14], 50 µM: 0.15 (n = 469); ci = [0.15, 0.15]) **[Fig. 4B]** In addition, the oROS-Gr green-to-red ratio predictably followed a sequence of exogenous H_2_O_2_ stimulation (100 uM) and DTT (10 mM) reduction **[Fig. 4C, Supp. Fig. 5]**.

**Figure 4.**
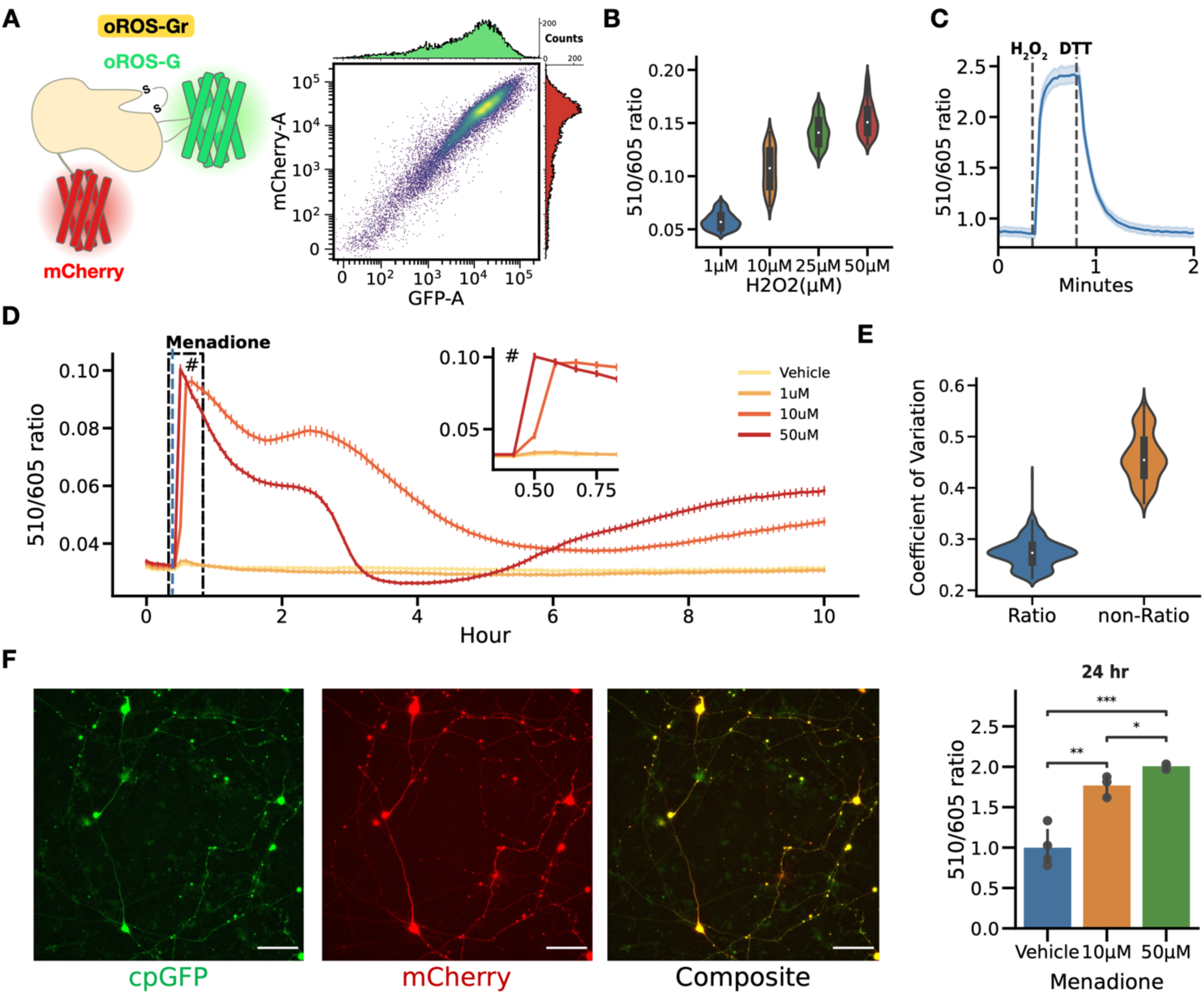
Ratiometric variant oROS-Gr improves temporal flexibility of H_2_O_2_ measurement. **A *left*** Schematic illustration of the oROS-Gr sensor design. oROS-Gr is an oROS-G variant with a C-terminus fusion of mCherry red fluorescent protein. ***right*** Flow cytometry result showed the linear relationship of green and red fluorescence levels of the oROS-G1r sensor expressed in HEK293 cells. **B** 510/605 nm emission ratio of oROS-G1r expressed in HEK293s stimulated with various concentrations of H_2_O_2_. Fluorescence emission ratios were captured a minute after the exposure. (n>100 cells per condition) **C** 510/605 nm emission ratio change of oROS-Gr during activation and reversal upon stimulation with 100µM H_2_O_2_ and 10mM DTT in HEK293 WT cells (n=82 cells). **D** Continuous monitoring of oROS-Gr fluorescence ratio for 10 hours in oROS-Gr stable cells in response to various concentrations of menadione (n>100 cells per condition). **E** Aggregated Coefficient of Variation (CoV) of 510/605 nm emission ratio and 510nm emission intensity from Figure 4D. Resting oROS-G1r ratio expressed in hiPSC-derived cortical neurons incubated in various levels of menadione for 24 hours. (n=3-4 wells per condition) **Descriptive statistics:** Error bars and bands represent the bootstrap confidence interval (95%) of the central tendency of values using the Seaborn (0.11.2) statistical plotting package. Cell-of-interests were collected from 3 biological replicates unless noted otherwise. **Inferential statistics:** F - t-test independent samples. *P < 0.05, **P < 0.01, ***P < 0.001.

Menadione has been widely used to model oxidative stress in biological systems. Still, studies to monitor its intracellular effect have been mostly limited to short time windows or non-continuous snapshots at varying time points. These studies did not provide insights into the real-time impact on redox homeostasis over a longer period. Here, we used to continuously measure (sampling every 5 minutes) the effects of menadione on cellular H_2_O_2_ levels over a ten-hour time window using stable oROS-Gr expressing HEK293 cells. Initially, menadione at 0, 1, 10, and 50 uM induced acute dose-dependent elevation of H_2_O_2_. However, within 30 minutes, the H_2_O_2_ levels at 10µM were higher than those at 50µM, which returned to a dose-dependent trend within four hours. **[Fig. 4D]** Further analysis of intracellular redox landscape analysis and functional role of putative cellular antioxidative elements^44,45^ is required to understand this phenomenon fully. Nevertheless, the temporally perplexing effects of menadione on intracellular H_2_O_2_ level gives us a cautionary insight to avoid the assumption that an oxidative agent such as menadione has always a direct dose-dependent effect on the intracellular peroxide level. We also tested if using oROS-Gr can improve the readout precision over oROS-G. We compared the coefficient of variation (CoV) of the ratiometric data (Em. 510/605) against data acquired in single wavelength mode (Em. 510) during long-term menadione exposure **[Fig. 4E].** The ratiometric readout showed about 2-fold lower CoV compared to the single wavelength mode, confirming improvements in precision [Ratiometric: 0.27 (n = 484), non-Ratiometric: 0.46 (n = 484)].

We confirmed the robust expression and functionality of oROS-Gr in various human stem cell-derived cells. For example, we measured peroxide levels in hiPSC-derived cortical neurons in response to 24-hour 10 µM and 50 µM Menadione incubation to be 1.77-fold and 2-fold of oROS-Gr ratio observed at vehicle negative control, respectively (Vehicle: 1.0; ci = [0.82, 1.21], 10 µM: 1.77; ci = [1.62, 1.88], 50 µM: 2.01; ci = [1.97, 2.03]) **[Fig. 4F]**. Next, we used the sarcoendoplasmic reticulum calcium ATPase (SERCA) blocker cyclopiazonic acid (CPA, 10 µM) to elevate Ca^2+^ in the cytosol of hiPSC-cardiomyocytes (CM)^46^. When hiPSC-CMs expressed oROS-Gr, we measured increased cytosolic peroxide levels within 2 hours of CPA incubation **[Supp. Fig. 4].** As previously reported, this confirms a tight coupling between intracellular Ca^2+^ and ROS levels^47–50^.

### Glucose-dependent basal oxidation level in mammalian cells

Superoxide and peroxide are continuously generated as byproducts through electron transfers during aerobic metabolism^51,52,53^. In this context, glucose, as one of the primary substrates of aerobic metabolic pathways, plays a crucial role in modulating cellular metabolic activity^54^. Intriguingly, low as well as high glucose levels were reported to result in depressed respiratory activity in cultured human podocytes^55^. The study also showed that the reduction of metabolic rates in high-glucose conditions can be reversed by incubation with the antioxidant NAC, indicating that respiratory suppression is correlated to oxidative stress. Thus, we hypothesized high glucose (HG = 25 mM) but also low glucose (LG = 1 mM) media would result in higher basal peroxide levels than medium glucose (MG = 10 mM). We incubated HEK293 cells for 48 hours in HG, NG, and LG media and compared the ratiometric oROS-Gr signals. Here, low and high glucose conditions caused higher peroxide levels than MG (MG: 0.38; ci = [0.378, 0.382], LG: 0.402; ci = [0.4, 0.404], HG: 0.392; ci = [0.389, 0.394]). **[Fig. 5A].** We directly measured metabolic activities and found that basal and maximum respiratory rates were also the lowest under low and high glucose conditions **[Fig. 5B, Supp. Fig. 7A, B]**, indicating an inverse correlation with increased peroxide levels. Indeed, cells that were pre-incubated with 1 mM of the antioxidant NAC under HG conditions brought the oROS-Gr level 84% closer to MG conditions, indicating modest suppression of oxidative stress **[Supp. Fig. 7C].**

**Figure 5.**
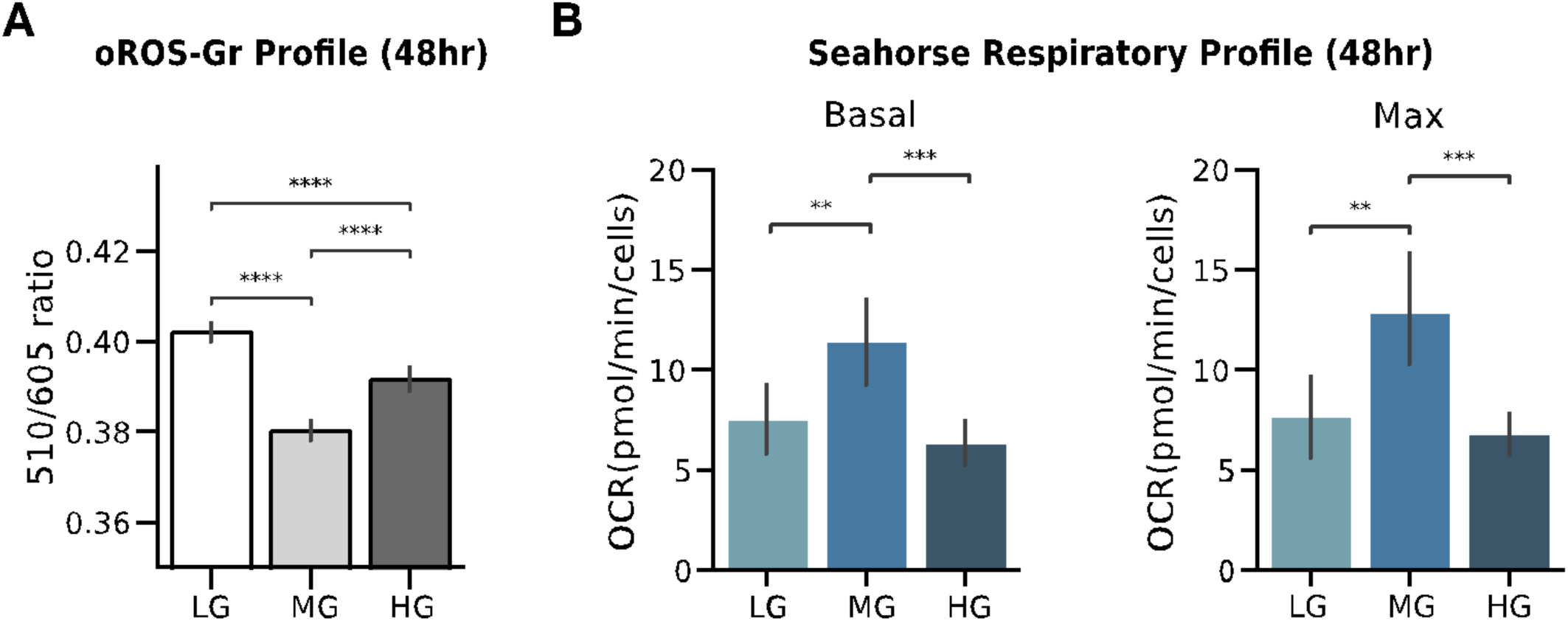
Glucose-dependent basal oxidation level in mammalian cells. **A** Observation of glucose-level dependent basal oxidation level of oROS-Gr (510/605 ratio) in oROS-Gr stable cells. The cells were incubated in LG (low glucose, 1mM), NG, and HG incubation for 48 hours after they were serum-deprived overnight. The sensors were least oxidized in NG, exhibiting a U-shape dose-dependent response (n>100 cells per condition). **B-C** LG and HG condition, which showed higher oxidation, also exhibits respiratory depression. **B** Basal respiratory level calculated from Seahorse assay, units in Oxygen Consumption Rate (pMol/min, n=24 wells per condition) **C** Max respiratory capacity calculated from Seahorse assay, units in Oxygen Consumption Rate (pMol/min, n=24 wells per condition). **Descriptive statistics:** Error bars and bands represent the bootstrap confidence interval (95%) of the central tendency of values using the Seaborn (0.11.2) statistical plotting package. Cell-of-interests were collected from 3 biological replicates unless noted otherwise. **Inferential statistics:** F - t-test independent samples. *P < 0.05, **P < 0.01, ***P < 0.001.

### Morphine elicits H_2_O_2_ generation in µ-opioid receptor-expressing neurons in the Ventral Tegmental Area *ex vivo*

We previously reported on a NADPH oxidase (NOX) dependent mechanism that inactivates opioid receptors in opioid receptor-expressing cell lines and *in vivo*^56,57^. Briefly, κ-opioid-receptor (KOR) and μ-opioid receptor (MOR) activation triggers cJUN N-terminal kinase (JNK) phosphorylation. Phosphorylated JNK then activates peroxiredoxin 6 (PRDX6), producing superoxide (SO) from NOX. SO can quickly oxidize the Gαi protein complex to inactivate the opioid receptors. This event can be captured using H_2_O_2_ as a marker of opioid receptor activation because superoxide is readily transformed into H_2_O_2_ by superoxide dismutase^58–61^ **[Fig. 6A]**. Using oROS-Gr, we tested whether morphine elicits transient peroxide generation in µ-opioid receptor-expressing neurons in the ventral tegmental area (VTA) of mice. The oROS signals were measured using 2-photon microscopy on *ex vivo* acute brain slice after viral delivery of the oROS-Gr gene (AAV1-DIO-oROS-Gr) into the VTA of MOR-Cre transgenic mouse. Expression of oROS-Gr in the VTA region was verified with one-photon confocal microscopy of post-mortem fixed brain slices **[Fig. 6B]**. 2-photon microscopy of ex vivo brain slice showed an acute increase of oROS-Gr response upon bath application of 1µM morphine over 30 minutes of monitoring which was blocked by the opioid receptor antagonist 1µM Naloxone (Normalized ΔoROS-Gr ratio to first 5 baseline frames, MOR: 3.35; ci = [1.94, 4.82], MOR+NLX: 0.63; ci = [0.42, 0.85]) **[Fig. 6C]**.

**Figure 6.**
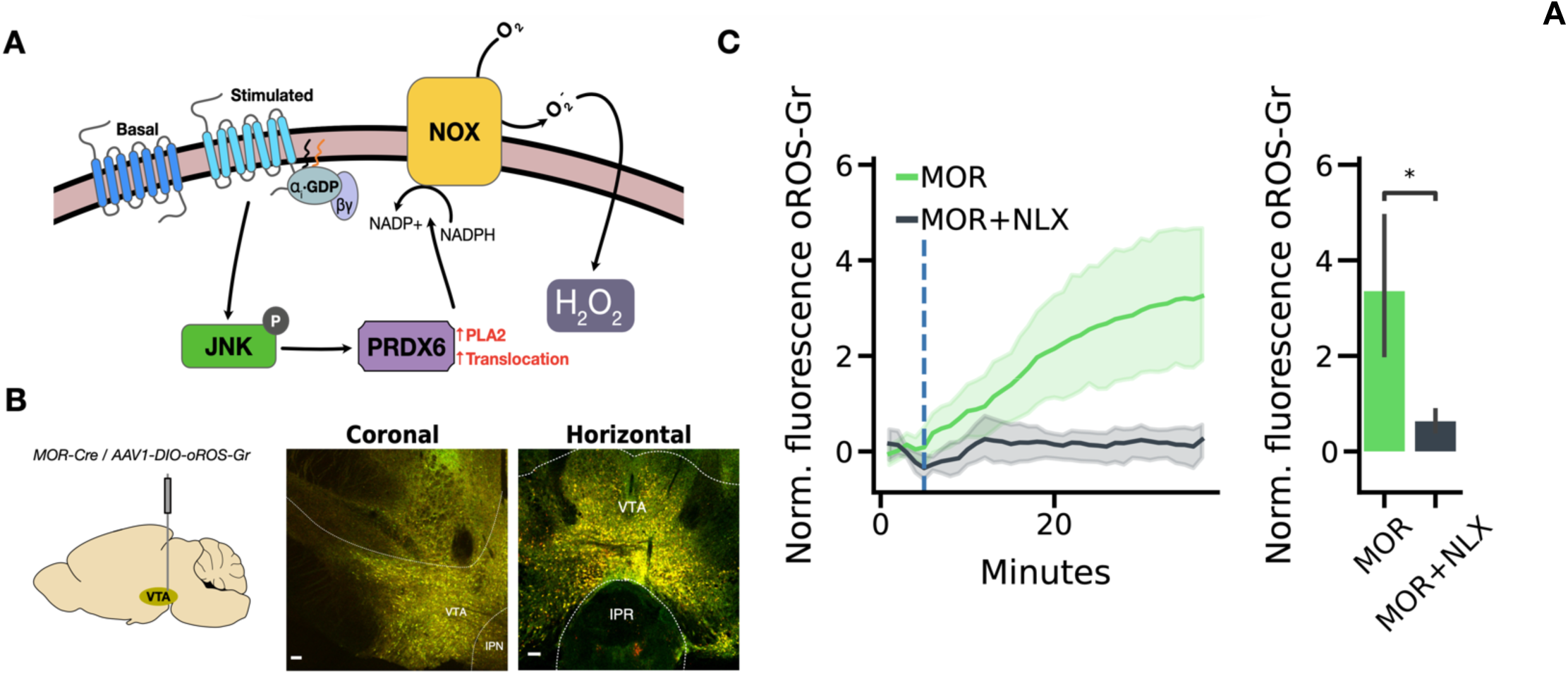
Morphine elicits H_2_O_2_ generation in µ-opioid receptor-expressing neurons in the Ventral Tegmental Area *ex vivo* and *in vivo*. A Schematic illustration of the JNK/PRDX6/NOX pathway for opioid receptor-dependent generation of H_2_O_2._ **B** Targeted expression of oROS-Gr in MOR positive neurons of Ventral tegmental Area (VTA) was achieved by AAV-DIO-oROSGr injection in VTA of MOR-Cre animals. Fluorescent images show histological validation of post-mortem confocal slice. **C** *Ex vivo* brain slice showing morphine-dependent H_2_O_2_ increase in µOR-positive neurons. (MOR: 1µM Morphine, n=10 slices. MOR + NLX: 1µM Morphine co-administered with 1µM Naloxone, n=9 slices) ***left*** real-time trace of morphine-induced oROS-Gr signals detected in µOR positive neurons of VTA. Naloxone, a competitive mu-opioid receptor antagonist, effectively diminishes Morphine-induced oROS-Gr signals. ***right*** maximal fluorescence response for each condition. **Descriptive statistics:** Error bars and bands represent the bootstrap confidence interval (95%) of the central tendency of values using the Seaborn (0.11.2). **Inferential statistics:** C - t-test independent samples. *P < 0.05, **P < 0.01, ***P < 0.001.

## Discussion

To further improve our understanding of redox biology, we need the ability to monitor oxidative agents in diverse, multi-faceted contexts. Our development of the oROS sensor framework represents a significant step in this direction. As a novel green fluorescent sensor, oROS-G demonstrates unparalleled sensitivity and response kinetics for H_2_O_2_ monitoring, surpassing the capabilities of previous ecOxyR-based sensors. This enhancement is largely attributed to our structural refinement of the sensor, where we relocated the cpGFP insertion site to maintain flexibility of C199-C208 loop of ecOxyR. Drawing inspiration from Akerboom et al., we incorporated bulky residues adjacent to the cpGFP barrel opening, exemplified by the E215Y mutation in oROS-G. Our study has revealed a novel insertion site within ecOxyR, paving the way for the creation of H_2_O_2_ sensors that are both fast and sensitive. Our hypothesis suggests that the flexible region in the ligand sensing domain is intrinsically linked to sensor function. These principles could lay the groundwork for future optogenetic sensors tailored to detecting other analytes.

Interestingly, the diversity of OxyR variants in nature, each characterized by a conserved peroxide oxidation mechanism, opens avenues for exploring a range of sensor functionalities. Notably, OxyRs from different bacterial strains exhibit distinct reduction mechanisms; ecOxyR predominantly follows a Grx (glutaredoxin)-dependent reduction pathway, where glutaredoxin proteins facilitate the reduction of oxidized proteins.^62^ In contrast, other variants like nmOxyR (*Neisseria meningitidis*) might employ a Trx (thioredoxin)-dependent reduction mechanism, involving the thioredoxin system known for mitigating cellular oxidative stress. (e.g. HyPer7)^41^ This variation necessitates further exploration of these domains for sensors in mammalian systems, where they could serve as complementary tools for dissecting peroxide biology in various redox environments.

Our study also demonstrated the practical versatility of oROS sensors in a range of experimental setups. With oROS-G, we successfully monitored H_2_O_2_ levels in astrocytes, both *in vitro* and *ex vivo*, shedding light on cellular redox states. Moreover, the ratiometric oROS-Gr sensor enabled us to observe the effects of glucose on cytoplasmic peroxide levels, which correlated with known patterns of mitochondrial oxidative stress.

Future studies should aim to clarify the sources of peroxide accumulation, considering factors like NADPH oxidase activity and mitochondrial respiration. Additionally, our work highlights the oROS sensor’s efficacy in detecting morphine-induced peroxide signals in µ-opioid receptor-positive neurons, further emphasizing its broad applicability.

In conclusion, the oROS sensors, exemplified by oROS-G and oROS-Gr, offer a new paradigm for studying peroxide biology. Their application across various model systems has the potential to revolutionize our approach to understanding and monitoring complex redox processes, with significant implications for unraveling the mechanisms underlying various oxidative stress-related diseases.

## Acknowledgments

J.D.L was supported by 1F31DA056121-01A1 and ISCRM Fellowship. A.B was supported by the Brain Research Foundation, UW Royalty Research Fund, UW ISCRM IPA, NIGMS R01 GM139850-01, P30 DA048736-01-Pilot, NIMH RF1MH130391, NINDS U01NS128537, NIDA R21DA051193 and the McKnight Foundation’s Technologies in Neuroscience Award. K. E was supported by T32AG066574. The research received additional support from the Lynn and Mike Garvey Imaging Core, the UW NAPE Center, and ISCRM Shared Equipment. We want to thank Dr. Randy Moon for his support. Also, this work was supported by the Institute for Basic Science, Center for Cognition and Sociaility (IBS-R001-D2) to C.J.L. We are also grateful to the IBS virus facility for providing a virus packaging service for in vivo experiments.

## Data Availability

Source data will be available via figshare shortly.

## Code Availability

Source code will be available at https://github.com/justindaholee/oROS-G_manuscript shortly.

## Ethics Statement

All animal procedures performed at the University of Washington have been approved by the University of 24 Washington’s Animal Use Committee (protocol #4422-01) and follow the National Institute of Health and the 25 Association for Assessment and Accreditation of Laboratory Animal Care International guidelines. Handling and animal care were at the Institute for Basic Science have been performed according to the Institutional Animal Care and Use Committee of the Institute for Basic Science.

## Material requests

### Plasmid name: Addgene #

AAV2_CAG_oROS-G_LF: 216116, AAV2_CAG_oROS-G : 216115 pCAG_oROS-Gr : 216114 pC3.1_CAG_oROS-Gr : 216113pC3.1_CMV_oROS-G_LF : 216112pC3.1_CMV_oROS-G : 216111

## Methods

### Protein structure analysis

Protein structure analysis and plotting were performed using Chimera-X-1.2.1. Oxidized [PDB:1I6A] and reduced [PDB:1I69] crystal structures of ecOxyR were imported from the Protein Data Bank (PDB). Pairwise residue distance between reduced and oxidized ecOxyR structure was achieved by aligning both structures using a matchmaker algorithm that superimposes protein structures by creating a pairwise sequence alignment and then fitting the aligned residue pairs to derive pairwise residue distances.

### Molecular Biology

oROS-HT variants were cloned based on the pC1 plasmid backbone from pC1-HyPer-Red (Addgene ID: 48249). Primers for point mutations or fragment assembly required to generate the oROS-HT screening variants were designed for In Vitro Assembly cloning (IVA) technique^63^, gibson assembly (New England Biolabs; E2611L) or blunt-end amplification for KLD-based site-directed mutagensis methods. Primers were ordered from Integrated DNA Technologies (IDT). All gene fragment amplifications were done using Seither Q5-polymerase (New England Biolabs; M0492L) or Superfi-II polymerase (Invitrogen; 12368010). Amplification of DNA fragments were verified with agarose gel electrophoresis. 30 minutes of DpnI enzyme treatment were done on every PCR product to remove the plasmid template from PCR samples. For IVA cloning circularization or assembly of the PCR products was achieved by transforming linear DNA products into competent E.Coli cells (DH5ɑ or TOP10) and grown on agar plates that contain either ampicillin or kanamycin selection antibiotic (50 µg/mL). For gibson assembly and KLD cloning, circularized DNA was transformed as above. Upon colony formation, single colonies were picked and grown in 5mL cultures containing LB Broth (Fisher BioReagents; BP9723-2) and selection antibiotic (ampicillin/kanamycin; 50 µg/mL) overnight (37°C, 230 RPM). DNA was isolated using Machery Nagel DNA prep kits (Machery Nagel; 740490.250). Sanger sequencing (Genewiz; Seattle, WA) or Whole plasmid nanopore sequencing (Plasmidsarus; Eugene, OR) of the isolated plasmid DNA was used to confirm the presence of the intended mutation. Genes encoding the final variants were cloned into a CAG-driven backbone, pCAG-Archon1-KGC-EGFP-ER2-WPRE (Addgene; #108423), using the methods above. England Biolabs; E2621L). All subsequences were verified with Sanger sequencing (Genewiz; Seattle, WA) or Whole plasmid nanopore sequencing (Plasmidsarus; Eugene, OR)

### Chemicals

H_2_O_2_ working solutions were freshly prepared before every experiment from H_2_O_2_ solution 30 % (w/w) in H_2_O (Sigma-Aldrich, H1009). Stock solution of Menadione (VENDOR, CAT) was prepared in 100% DMSO at 50mM. Stock solution of Cyclopiazonic Acid (Tocris, 1235) was prepared in 100% DMSO at 20mM. Chemicals specific to other method sections can be found in their respective sections.

### Cell culture and transfection

Human Embryonic Kidney (HEK293; ATCC Ref: CRL-1573) cells were cultured in Dulbecco’s Modified Eagle Medium + GlutaMAX (Gibco; 10569-010) supplemented with 10% fetal bovine serum (Biowest; S1620). When cultures reached 85% confluency, the cultures were seeded at 150,000/75,000 cells per well in 24/48-well plates, respectively. 24 hours after cell seeding, the cells were transfected using Lipofectamine3000 (Invitrogen; L3000015) at 1000/500 ng of DNA per well of a 24/48-well plate, according to the manufacturer’s instructions.

### Primary rat neuron isolation

Primary cortical neurons were prepared as previously described^47,48^. Briefly, 24-well tissue culture plates were coated with Matrigel (mixed 1:20 in cold-PBS, Corning; 356231) solution and incubated at 4°C overnight prior to use. Sterile dissection tools were used to isolate cortical brain tissue from P0 rat pups (male and female). Tissue was minced until 1mm pieces remained, then lysed in equilibrated (37°C, 5% CO_2_) enzyme (20 U/mL Papain (Worthington Biochemical Corp; LK003176) in 5mL of EBSS (Sigma; E3024)) solution for 30 minutes at 37°C, 5% CO_2_ humidified incubator. Lysed cells were centrifuged at 200xg for 5 minutes at room temperature, and the supernatant was removed before cells were resuspended in 3 mLs of EBSS (Sigma; E3024). Cells were triturated 24x with a pulled Pasteur pipette in EBSS until homogenous. EBSS was added until the sample volume reached 10 mLs prior to spinning at 0.7 rcf for 5 minutes at room temperature. Supernatant was removed, and enzymatic dissociation was stopped by resuspending cells in 5 mLs EBSS (Sigma; E3024) + final concentration of 10 mM HEPES Buffer (Fisher; BP299-100) + trypsin inhibitor soybean (1 mg/ml in EBSS at a final concentration of 0.2%; Sigma, T9253) + 60 µl of fetal bovine serum (Biowest; S1620) + 30 µl 100 U/mL DNase1 (Sigma;11284932001). Cells were washed 2x by spinning at 0.7 rcf for 5 minutes at room temperature and removing supernatant + resuspending in 10 mLs of Neuronal Basal Media (Invitrogen; 10888022) supplemented with B27 (Invitrogen; 17504044) and glutamine (Invitrogen; 35050061) (NBA++).

After final wash spin and supernatant removal, cells were resuspended in 10 mLs of NBA++ prior to counting. Just before neurons were plated, matrigel was aspirated from the wells. Neurons were plated on the prepared culture plates at desired seeding density. Twenty-four hours after plating, 1µM AraC (Sigma; C6645) was added to the NBA++ growth media to prevent the growth of glial cells.Plates were incubated at 37°C and 5% CO_2_ and maintained by exchanging half of the media volume for each well with fresh, warmed Neuronal Basal Media (Invitrogen; 10888022) supplemented with B27 (Invitrogen; 17504044) and glutamine (Invitrogen; 35050061) every three days.

### Human primary astrocytes, and stem cell derived cardiomyocytes and neurons

Astrocytes: Human primary cortical astrocytes were purchased from ScienCell Research Laboratories (Carlsbad, CA) and were stored, thawed and sub-cultured based on the manufacturer’s protocol. Briefly, the astrocytes were cultured for 72 h in a base medium with an astrocyte growth supplement and fetal bovine serum provided by the same manufacturer. Cultures were maintained in a 37°C/5% CO_2_ incubator throughout the culture period, and the astrocytes with low passage numbers (p0-p3) were used to guarantee consistent phenotype expression. When the culture became 70% confluent, the cells were dissociated with TrypLE (Thermo Fisher), followed by passaging on the PDL-coated 24 cover glasses for oROS-G1 transfection. The transfected cells were then cultured for an additional 96 h before H_2_O_2_ treatment (10 µM, 100 µM) for recording the fluorescence response upon H_2_O_2_ stimulation.

Cardiomyocytes: Undifferentiated IMR90 (WiCell) hiPSCs were maintained on Matrigel (Corning) coated tissue culture plates in mTeSR1 (Stemcell Technologies). Cardiomyocyte directed differentiation was performed using a modified small molecule Wnt-modulating protocol using Chiron 99021 and IWP-4 as previously described.^64,65^. Lactate enrichment was performed following differentiation to purify hiPSC-CMs.^66^

Cortical neurons: Neurons were generated from the previously characterized wild type CV background human induced pluripotent stem cell line (Young et al. 2015). Neural progenitor cells (NPCs) from this cell line were differentiated from hiPSCs using dual-SMAD inhibition and NPCs were differentiated to neurons as previously described (Knupp et al., 2020; Shin et al., 2023). Briefly, for cortical neuron differentiation from NPCs, NPCs were expanded into 10 cm plates in Basal Neural Maintenance Media (BNMM) (1:1 DMEM/F12 (#11039047 Life Technologies) + glutamine media/neurobasal media (#21103049, GIBCO), 0.5% N2 supplement (# 17502-048; Thermo Fisher Scientific,) 1% B27 supplement (# 17504-044; Thermo Fisher Scientific), 0.5% GlutaMax (# 35050061; Thermo Fisher Scientific), 0.5% insulin-transferrin-selenium (#41400045; Thermo Fisher Scientific), 0.5% NEAA (# 11140050; Thermo Fisher Scientific), 0.2% β-mercaptoethanol (#21985023, Life Technologies) + 20 ng/mL FGF (R&D Systems, Minneapolis, MN). Once the NPCs reached 100% confluence, they were switched to Neural Differentiation Media (BNMM +0.2 mg/mL brain-derived neurotrophic factor (CC# 450–02; PeproTech) + 0.2 mg/mL glial-cell-derived neurotrophic factor (CC# 450–10; PeproTech) + 0.5 M dbcAMP (CC# D0260; Sigma Aldrich). Neural Differentiation Media was changed twice a week for 21 days at which point the differentiation is considered finished. Neurons were replated at a density of 500,000 cells/cm^2^.

### Imaging

Imaging experiments described in this study were performed as follows unless specifically noted. Epifluorescence imaging experiments were performed on a Leica DMI8 microscope (Semrock bandpass filter: GFP ex/em: FF01-474-27/FF01-520-35, RFP ex/em:FF01-578-21/FF01-600-37) controlled by MetaMorph Imaging software, using a sCMOS camera (Photometrics Prime95B) and 20x magnification lens (Leica HCX PL FLUOTAR L 20x/0.40 NA CORR) or 10× objective (Leica HCX PL FLUOTAR L 10x/0.32 NA). Confocal imaging experiments were performed on a Leica SP8 confocal microscope from the Lynn and Mike Garvey Imaging Core at the Institute of Stem Cell and Regenerative Medicine. Cells were imaged in live cell imaging solution with 10mM glucose (LCIS+, Gibco, A14291DJ). Image analysis methods described below.

### Analysis

Analysis of cell fluorescence imaging data was done by FUSE, a custom cloud-based semi-automated time series fluorescence data analysis platform written in Python. First, the cell segmentation quality of the selected Cellpose^81^ model was manually verified. For the segmentation of cells expressing cytosolic fluorescent indicators, model ‘cyto’ was selected as our base model. If the selected Cellpose model was low-performing, we further trained the Cellpose model using the Cellpose 2.0 human-in-the-loop system^82^. Using an “optimized” segmentation model, fluorescence time-series data is extracted for each region of interest. This allows for unbiased extraction of change in cellular fluorescence information for a complete set of experimental samples. Extracted fluorescence data is normalized as specified in the text using custom python script.

### Astrocyte study

#### Primary mouse astrocyte culture

Primary mouse cultured astrocytes were prepared from P1-P3 C57BL/6J mouse pups as previously described.^67^ Briefly, 60-mm culture dishes were coated with 0.1 mg/ml poly-D-lysine (PDL, Sigma; P6407) solution prior to use. The hippocampal tissue was isolated, and dissociated into single cell suspension by tituration in Dulbecco’s modified Eagle’s meidum supplemented with 4.5 g/L glucose, L-glutamine, sodium pyruvated (DMEM, Corning; 10-013-CV) + 10% heat-inactivated horse serum (Gibco; 26050-088) + 10% heat-inactivated fetal bovine serum (Dawin bio; A0100-010) + 1000 unit/ml penicillin-streptomycin (Gibco; 15140122). Dissociated cells were plated onto the PDL coated dishes. Cultures were maintained at 37°C in a humidifed atmosphere cotanining 5% CO_2_ incubator. On the third day, cells were vigorously washed with repeated pipetting using medium to get rid of debris and other floating cell types.

On the 10th day of culture, cultured primary astrocytes were electroporetically transfected with oROS-G plasmid with a voltage protocol (1200 V, 20 pulse width, 2 pulses) using the Microporator (Invitrogen Neon Transfection System; MPK5000S) and replated onto converglass coated with PDL (Sigma; P6407) or µ-Plate 96 Well Black (ibidi; 89626).

#### Imaging of cultured primary mouse astrocytes

On the 14th day of culture, the oROS-G transfected cultured primary astrocytes were transferred to a recording chamber which were mounted on an inverted Nikon Ti2-U microscope and continuously perfused with an external solution contained (in mM): 150 NaCl, 10 HEPES, 5.5 glucose, 3 KCl, 2MgCl2, 2 CaCl2, and pH adjusted to pH 7.3. Intensity images of 525 nm wavelength were taken at 485 nm excitation wavelengths using ORCA-Flash4.0 CMOS camera (Hamamatsu; C13440). Imaging workbench (INDEC Biosystem) and ImageJ (NIH) were utilized for image acquisition and ROI analysis of cultured astrocytes. To examine H_2_O_2_-dose dependent responses of oROS-G transfected cultured astrocytes, concentration of 10 and 100 µM of H_2_O_2_ (Sigma; 88597) were introduced by bath application. The peak response of the sensor was normalized to its baseline (ΔF/Fo), wich was measured 90 s before the introducing H_2_O_2_. For confocal live-cell imaging and monitoring antioxidant drugs, confocal imaging was performed by using Nikon A1R confocal microscope mounted onto a Nikon Eclipse Ti body with 20x objective lens. A Live-cell imaging chamber and incubation system were used for maintaining environmental condtions at 10% CO2 and 37°C during 40 h continuous recordings. Images were acquired by using NIS-element AR (Nikon). For image analysis, NIS-element (Nikon) and ImageJ (NIH) were used.

#### Animals

All APP/PS1 mice were group-housed in a temperature- and humidity-controlled environment with a 12 h light/dark cycle and had a free access to food and water. All animal care and handling was approved by the Institutional Animal Care and Use Committee of the Institute for Basic Science (Daejeon, Korea).

#### Virus injection

The AAV5-GFAP104-oROS-G viral vector was cloned and AAV containing GFAP-104-oROS-G was packaged by the IBS virus facility (Daejeon, Korea). Mice were deeply anesthetized via vaporized 1% isoflurane and immobilized in a stereotaxic (RWD Life Science). Following an incision on the midline of the scalp, bilateral craniotomies were performed above the hippocampus CA1 (anterior/posterior, -2 mm; medial/lateral, ±1.6 mm; dorsal/ventral, -1.45 mm from the bregma) using a microdrill. The virus was bilaterally microinjected (0.1 μl/min for 10 min; total 0.8 μl) using a syringe pump (KD Scientific).

#### oROS-G imaing of GFAP-positive astrocytes in the brain slices

A total of 2 weeks after the virus injection into the hippocampus, animals were anesthetized with vaporized 1% isoflurane and decapicated. The brains were submerged in chilled cutting solution that contained (in mM): 250 Sucrose, 26 NaHCO3, 10 D(+)-gluocse, 4 MgCl2, 0.1 CaCl2, 2.5 KCl, 2 Sodium Pyruvate, 1.25 NaH2PO4, 0.5 ascrobic acid, and pH adjusted to pH 7.4. Coronal slices (300 μm thick) were prepared with a vibrating-knife microtome D.S.K LinearSlicer pro 7 (Dosaka EM Co. Ltd). For stabilization, brain slices were incubated at room temparature for at least 1 h before imaging. For imaging, the slices were transferred to a recording slice chamber which were mounted on an upright Zeiss Examiner D1 microscope and continuously perfused with an artificial cerebrospinal fluid (aCSF) solution that contained (in mM): 130 NaCl, 24 NaHCO3, 1.25 NaH2PO4, 3.5 KCl, 1.5 MgCl2, 1.5 CaCl2, D(+)-glucose, and pH adjusted to pH 7.4. All solutions were equilibrated with 95% O2 and 5% CO2. Imaging was acquired at 0.25 frame per second with 60X water-immersion objective lens, a ORCA-Flash4.0 CMOS camera (Hamamatsu; C13440), and a LED (CoolLED) filtered with 485-nm fluorescence was applied. Imaging workbench (INDEC Biosystem) and ImageJ (NIH) were utilized for image acquisition and ROI analysis. To examine H_2_O_2_-dose dependent responses of sensor-expressing astrocytes, concentration of 10 and 100 µM of H_2_O_2_ were introduced by bath application. The peak response of the sensor was normalized to its baseline (ΔF/Fo), wich was measured 90 s before the introducing H_2_O_2_. To measure endogenous H_2_O_2_ in astrocytes of APP/PS1 mice and their littermates, we used 10 mM DTT (Thermo; R0861). This method reduced the oROS-G sensor bound to H_2_O_2_, resulting in fluorescence below the baseline levels. These reduced fluorescence responses were normalized to its baseline (ΔF/Fo), suggesting the basal endogenous H_2_O_2_ levels.

### Generation of stable oROS-Gr expressing HEK293 cells

HEK293 cells in a T75 flask were transfected (using lipofection, as described above) with oROS-Gr-P2A-Puromycin plasmid. 3 Days after the transfection, cells were passaged to 2 T75 flasks. 2 Days after, puromycin-based selection was performed for a week using complete DMEM media (as previously described) supplemented with puromyocin (1µg/mL). Cells after selection were expanded for 3 passages. Enrichment of cell population with robust oROS-Gr expression were achieved with BD FACSAria II Cell Sorter at Flow and Imaging Core Lab of University of Washington South Lake Union Campus.

### Glucose experiment and Seahorse Assay

oROS-Gr stable cells cultured in complete DMEM with 10mM glucose were plated at 75,000/well in 24-well plates. oROS-Gr stable cells were plated at 75,000/well in 24-well plates. 1 day post seeding, FBS in the DMEM media was brought down to 2% from 10%. 2 day post seeding cells were in serum-free DMEM with varying level of glucose. Mannose was supplemented as needed to keep osmotic pressure of each media consistant (final total sugar content: 25mM). oROS-Gr ratio (GFP/RFP) were imaged in LCIS media with varying glucose and mannose level. For Seahorese assay, oROS-Gr stable cells mentioned above were plated in a Matrigel-coated 96 well Seahorse plate at a density of 2 × 10^5^ cells/well for an equivalent procedure as above. The MitoStress protocol in the Seahorse XF96 Flux Analyzer (Agilent Technologies, Santa Clara, CA, USA) was performed two weeks later. An hour before the assay, the culture media was replaced with base media (Agilent Seahorse XF base medium, 103334-100 Agilent Technologies, Santa Clara, CA, USA) supplemented with 25 mM glucose and 1 mM Sodium pyruvate (11360070 Gibco/Thermo Scientific, Waltham, MA, USA). Substrates and select inhibitors of the different complexes were injected during the measurement to achieve final concentrations of oligomycin (2.5 μM), FCCP (1 μM), rotenone (2.5 μM) and antimycin (2.5 μM). The oxygen consumption rate (OCR) values were then normalized with readings from Hoechst staining (HO33342 Sigma-Aldrich, St. Louis, MO, USA), which corresponded to the number of cells in the well.

### µ-opioid receptor study

#### AAV

An adenovirus associated double floxed inverted (AAV1-DIO) virus was generated containing the oROS-Gr by cloning oROS-Gr into pAAV1-Ef1a-DIO using Nhe1 and Asc1 restriction sites. AAV1 were prepared by the NAPE Molecular Genetics Resources Core as described previously (Gore, et al, 2013). HEK293T cells were transfected with 25 μg AAV1 vector plasmid and 50 μg packaging vector (pDG1) per 15 cm plate. Two days after transfection, cells were harvested and subjected to three freeze–thaw cycles. The supernatant was transferred to a Beckman tube containing a 40% sucrose cushion and spun at 27,000 rpm overnight at 4°C. Pellets were resuspended in CsCl at a density of 1.37 g/ml and spun at 65000 rpm 4 hours at 4°C. 1 ml CsCl fractions were run on an agarose gel, and genome-containing fractions were selected and spun at 50000 rpm overnight at 4°C. The 1 ml fractions were collected again, and genome containing fractions were dialyzed overnight. The filtered solution was transferred to a Beckman tube containing a 40% sucrose cushion and spun at 27,000 rpm overnight at 4°C. The pellet (containing purified AAV) was resuspended in 150 μl 1× HBSS. Virus was aliquoted and stored at -80 ° C until use.

#### Animals and surgeries

Test naive C57BL/6 mice are ear punched at least 21 days after birth and genotyped using Transnetyx genotyping services. PCR screening was performed for the presence of Cre recombinase. For slice acquisition, mice ideally between 5-7 weeks of age are Injected with 0.5uL AAV1-Efla-FLEX-oROS-mCherry (CITE) construct containing oROS-Gr into a MOR CRE positive mouse bilaterally into the VTA using coordinates: ML: +/- 0.5, AP: -3.28, DN: -4.5 zeroed at bregma. Isoflurane is used for anesthesia and carprofen for pain relief. Mount on a stereotaxic alignment system and use a Hamilton 2.0uL model 7002 KH syringe for injections. Similarly, for fiber photometry experiments, mice are injected with 0.5uL AAV1-Efla-FLEX-oROS-mCherry unilaterally at a 15 degree angle, using the coordinates ML: -1.71, AP: -3.28, DN: -4.64 then implanted with a 400/430 uM diameter Mono fiberoptic cannula from Doric Lenses.

#### 2-photon imaging of µ-opioid receptor expressing neurons in VTA

Wait two-four weeks before removing the brain and obtaining 200um horizontal slices using a vibratome. Use NMDG as a cutting solution (92mM NMDG, 2.5mM KCl, 1.25mM NaH2PO4, 30mM NaHCO3, 20mM HEPES, 25mM Glucose, 2mM Thiourea, 5mM Na-ascorbate, 3mM Na-pyruvate, pH to 7.4, 0.5mM CaClx4H2O, 10mM MgSO4x7H2O). Place in an NMDG solution at 32 C for 15 minutes. Afterwards allow slices to incubate for an hour in a HEPES solution (92mM NaCl, 2.5mM KCl, 1.25mM NaH2PO4, 30mM NaHCO3, 20mM HEPES, 25mM Glucose, 2mM Thiourea, 5mM Na-Ascorbate, 3mM Na-Pyruvate). Begin image collection using a Bruker Investigator 2-photon microscope, software Prairie View 5.5, simultaneously collecting both the mCherry (1040nM fixed) and GFP (920nM tunable) signals with a Nikon 16X water immersion objective, as well as a z-stack spanning 60um across an hour time course. During this collection time establish a baseline for 7 minutes by washing on with ACSF buffer solution (124mM NaCl, 3mM KCl, 2mM MgSO4, 1.25mM NaH2PO4, 2.5mM CaCl2, 26mM NaHCO3, 10mM Glucose) at 32C, then wash on the treatment (10uM morphine or if treating with 10uM morphine and 1uM naloxone wash on naloxone for an additional 7 minutes before treatment) for 30 minutes. For coronal images, the animal was perfused intracardially with phosphate-buffered saline (PBS) and 10% formalin. Brains were stored in 10% formalin for up to 24 hours then switched to a 20% sucrose solution at 4C until sectioning. Coronal slices of the VTA were collected at 40uM each and mounted using VECTASHIELD HardSet mounting Medium with DAPI. No antibody was applied. Horizontal images were collected using slices that underwent 2P imaging and were put into 10% formalin. Confocal images are taken with the Leica SP8x Confocal microscope located in the Keck Center at UW.

## Notes

### Competing Interest Statement

The authors have declared no competing interest.

