## Supplemental Figures and tables for "Structure-guided engineering of a fast genetically encoded sensor for real-time H_2_O_2_ monitoring"

**Supplementary Figure 1** Structure-guided engineering of oROS-G

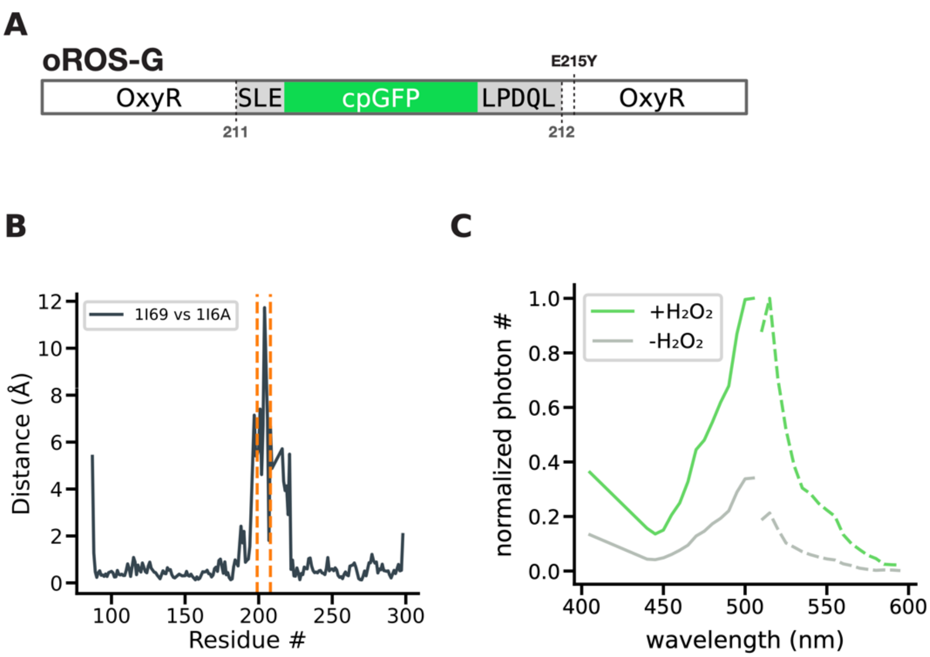


**A** Schematic description of oROS-G sequence design **B** Reduced [PDB:1I69] and oxidized [PDB:1I6A] form of OxyR was superimposed and calculated to residue-to-residue distance to estimate the degree of conformational change. Gray lines indicate C199 and C208. **C** Excitation (solid line) and Emission (dotted line) spectra of oROS-G expressed in HEK293s acquired with Leica Stellaris confocal microscope.

**Supplementary Figure 2** Spatiotemporal peroxide diffusion monitoring with oROS-G.

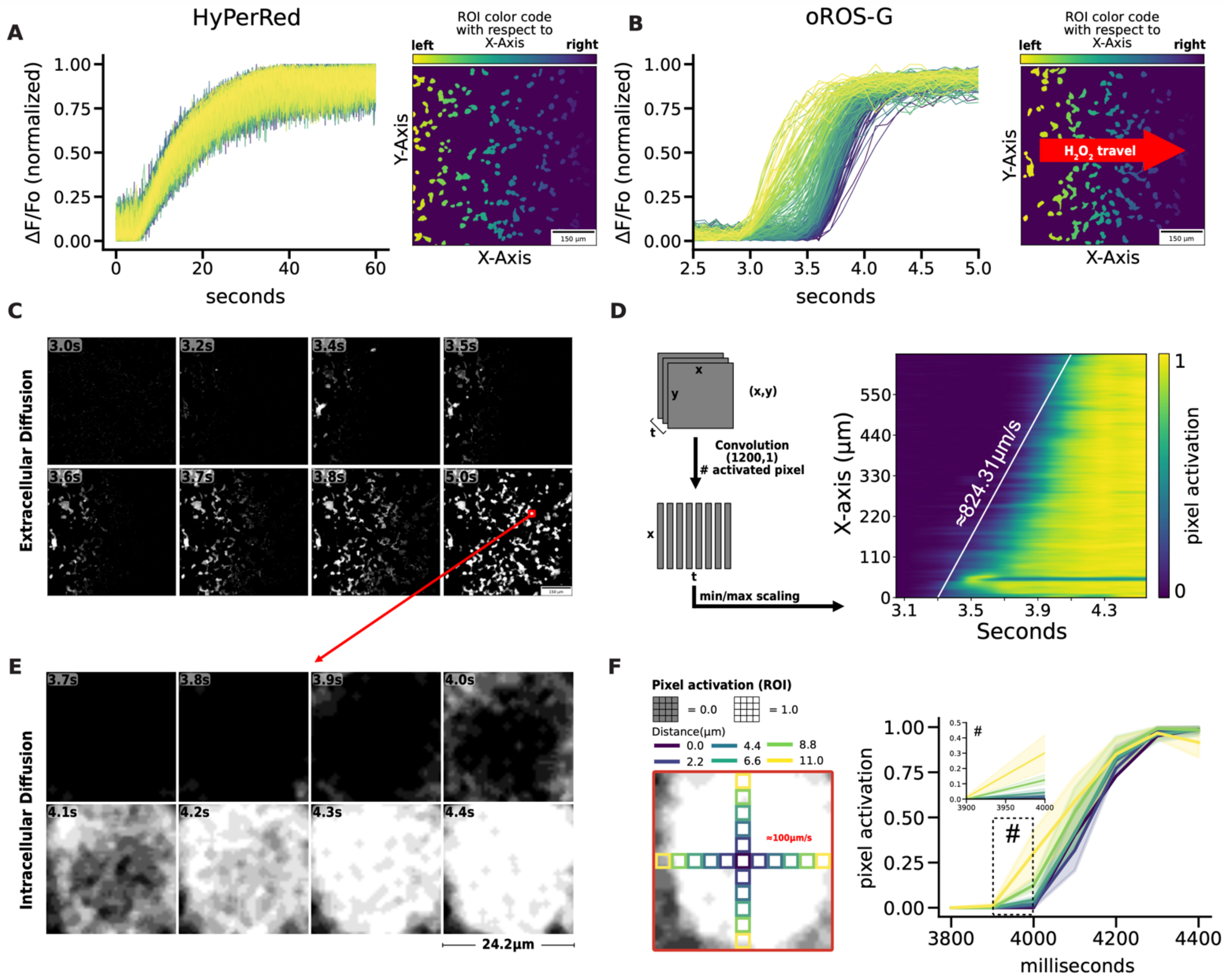


**A-B** Fluorescence response of oROS-G and HyPerRed to 300μM $H_2O_2$ stimulation captured at 10Hz. Region-of-interests (ROI) were color-coded for their respective X-axis location. **A** The stimulation induces uniform population-level activation in HyPerRed. **B** In contrast, stimulation induces spatially defined temporal gradient oROS-G activation that captures peroxide travel across the field of view. **C** Representative time-series frames from **Supp. Fig 2B,** Pixel values are gated to be “Black (0)” at 0~50% sensor activation and White (1) at 50%~100% sensor activation to enhance visualization. **D** 2D visualization of spatiotemporal oROS-G activation. Each time frame ('t') is represented by compressing the activated pixel data along the y-axis onto the x-axis. This process transforms each frame's data into a one-dimensional array of 1200 elements which then was min/max scaled to reflect the degree of pixel activation at each x-axis point. From the representation, peroxide travel speed during the media mixing is observed to be ≈824μm/s. **E** Zoomed-in representative time-series frames showing intracellular diffusion of peroxide. **F** Regions of Interest (ROIs) with a size of 3x3 pixels were organized radially and categorized by their proximity to the cell's center, indicated through color coding. The pixel activation levels within these ROIs were then plotted accordingly.

**Supplementary Figure 3** Characterization of ultrasensitive and fast peroxide sensor, oROS-G.

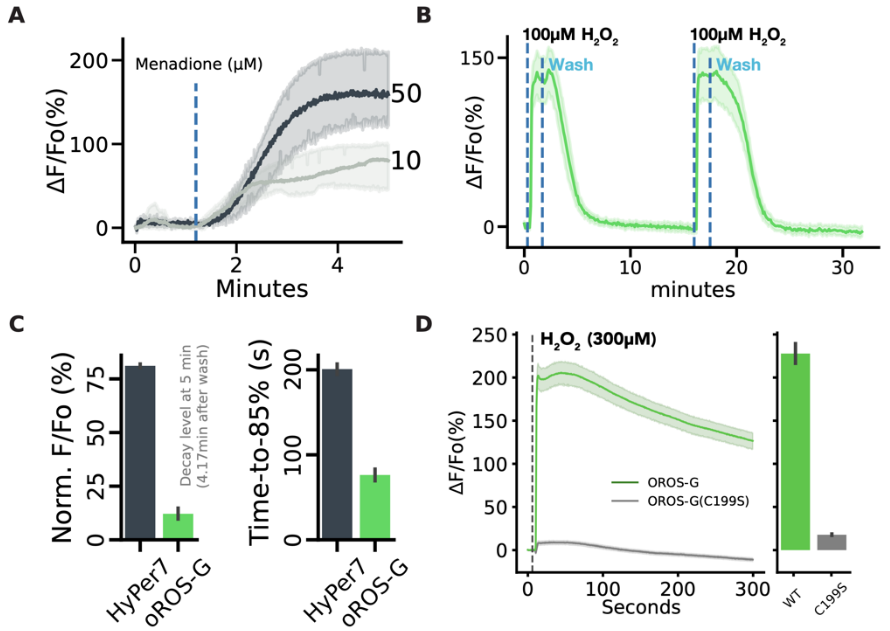


**A** Dose-dependent fluorescence response of oROS-G to high (50μM) or low (10μM) menadione expressed in human primary astrocytes. (n=3 cells per condition) **B** HEK293s expressing oROS-G were stimulated with 100μM $H_2O_2$ followed by media wash. The sequence was repeated twice. **C** derived *from Fig. 2F:* (a) Normalized ΔF/Fo (%) level of oROS-G and HyPer7 at 4.17 minutes after wash-induced reduction. The smaller value indicates a more efficient reduction. (b) Time measured for reduction to 85% fluorescence from the sensor saturation. **D** HEK293s expressing oROS-G and oROS-G1-C199S (loss-of-function) were stimulated with 300μM $H_2O_2$. The barplot represents the mean of the maximum fluorescent responses. **Descriptive statistics:** Error bars and bands represent the bootstrap confidence interval (95%) of the central tendency of values using the Seaborn (0.11.2) statistical plotting package. Cell-of-interests were collected from 3 biological replicates unless noted otherwise.

**Supplementary Figure 5**

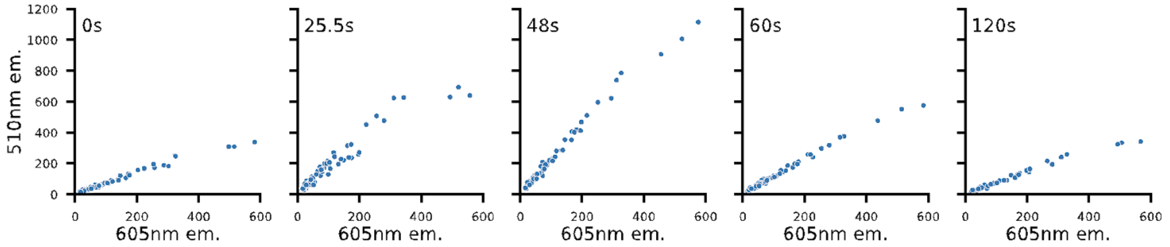


Distribution of 510 nm and 605 nm emission intensity of each ROI at selected temporal snapshots from Figure 4C.

**Supplementary Figure 6**

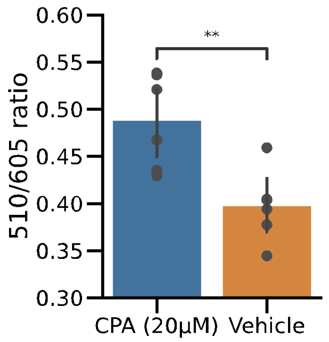


Resting oROS-Gr ratio expressed in hiPSC-derived cardiomyocytes incubated in SERCA blocker cyclopiazonic acid (CPA) for 2 hours. (CPA: 0.49 (n = 6); ci = [0.45, 0.52], Vehicle: 0.4 (n = 6); ci = [0.37, 0.43]) **Descriptive Statistics:** Error bars and bands represent the bootstrap confidence interval (95%) of the central tendency of values. **Inferential Statistics:** t-test independent samples. *P < 0.05, **P < 0.01, ***P < 0.001.

**Supplementary Figure 7**

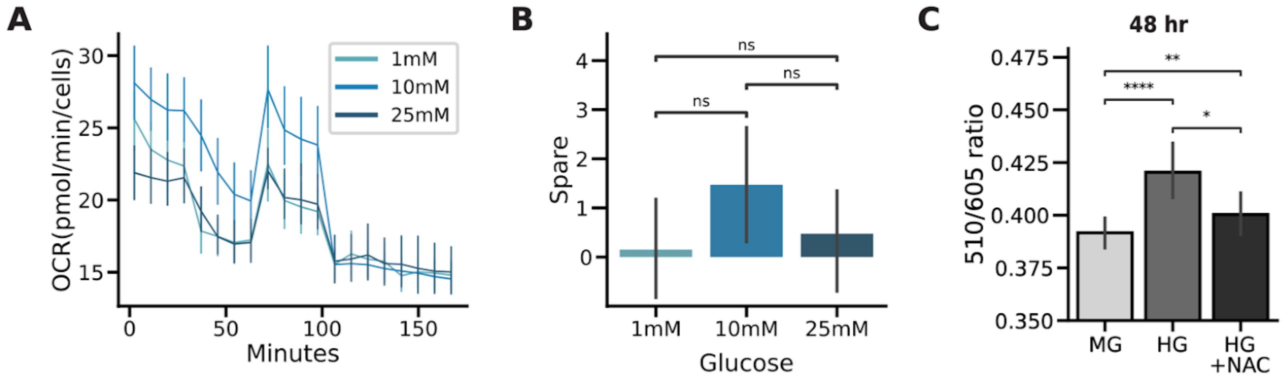


**A** Seahorse assay results after incubating oROS-Gr stable HEK293 cells in various glucose conditions (n=24 wells per condition). **B** Spare respiratory capacity calculated from Seahorse assay, units in Oxygen Consumption Rate (pMol/min) **C** Observation of glucose-level dependent basal oxidation level of oROS-Gr (510/605 ratio) in stably expressing HEK293 cells. The cells were incubated in MG (Medium glucose, 10mM), HG (high glucose, 25mM), or HG+NAC (negative control, HG +1mM of NAC) for 48 hours after they were serum-deprived overnight. NAC diminishes the oxidation level of HG (n>100 cells per condition). **Descriptive statistics:** Error bars and bands represent the bootstrap confidence interval (95%) of the central tendency of values using the Seaborn (0.11.2) statistical plotting package. Cell-of-interests were collected from 3 biological replicates unless noted otherwise. **Inferential statistics:** F - t-test independent samples. *P < 0.05, **P < 0.01, ***P < 0.001.
